# *Draft genome sequence of Stenotrophomonas goyi* sp. nov., a novel bacterium associated with the alga *Chlamydomonas reinhardtii*

**DOI:** 10.1101/2023.05.04.539380

**Authors:** María Jesus Torres, Neda Fakhimi, Alexandra Dubini, David González-Ballester

## Abstract

*Stenotrophomonas goyi* sp. nov. has been isolated from a contaminated algal culture (*Chlamydomonas reinhardtii*). Its genome has been fully sequenced (4,487,389 base pairs) and a tentative annotation is provided (4,147 genes). The genome information suggests that *S. goyi* sp. nov. is unable to use sulfate and nitrate as sulfur and nitrogen sources, respectively. Growth tests have confirmed the dependence of the sulfur-containing amino acids methionine and cysteine. The potential biotechnological interest of this bacteria is discussed here and in a related research paper (Fakhimi et al., 2023b).

## INTRODUCTION

The first described specie of the Stenotrophomonas genus was S. *maltophilia* which was a Gram-negative bacterium originally named as *Pseudomonas maltophilia*, and later transferred in 1993 to the new genus Stenotrophomonas, which was composed solely by *S. maltophilia*. In 2001 this specie was moved to the genus Xanthomonas before it was finally moved back again in 2017 to its own genus when *Stenotrophomonas pictorum* was identified (Ryan et al., 2009; Wei et al., 2021). Currently, Stenotrophomonas is a genus comprising at least 19 validated species (https://lpsn.dsmz.de/genus/stenotrophomonas) (Parte et al., 2020). However, the molecular taxonomy of the genus is still somewhat unclear, and all its members are considered as “orphan species”. All Stenotrophomonas ssp. shown intraspecific heterogeneity with high phenotypic, metabolic, and ecological diversity (Ryan et al., 2009).

The main reservoirs of Stenotrophomonas spp. are soil and plants, although they are ubiquitously present in different environments, including opportunistic human pathogens such as *S. maltophilia* (Ryan et al., 2009).

Stenotrophomonas spp. have promising potential for different biotechnological applications. Some Stenotrophomonas spp. are of interest in agriculture due to their ability to promote growth in different plant species, some of which are even capable of establishing symbiotic relationships with plants. This plant growth promotion is related to the capacity of some Stenotrophomonas spp. to produce the plant growth hormone indole⍰3⍰acetic acid (IAA), fix nitrogen, oxidate elemental sulfur (S) to sulfate, or biocontrol plant pathogens (Banerjee and Yesmin, 2008; Park et al., 2005; Ryan et al., 2009; Suckstorff and Berg, 2003).

Moreover, they are also considered good candidates for bioremediation due to their tolerance to heavy metals and capability to metabolize a large variety of organic molecules, including phenolic and aromatic compounds (Liu et al., 2007; Mora-Salguero et al., 2019; Pages et al., 2008; Ryan et al., 2009). Finally, some Stenotrophomonas spp. can synthetize useful bioproducts such as antimicrobial and enzymes of biotechnological interest (Rivas-Garcia et al., 2022; Wolf et al., 2002).

Here we report the genome of *Stenotrophomonas goyi*. sp. nov. isolated from a contaminated microalgae (*Chlamydomonas reinhardtii*) culture. This alga culture was simultaneously contaminated with *S. goyi, Microbacterium fakhimi* (Fakhimi et al., 2023a) and *Bacillus cereus*. The metabolic interactions established between these four microorganisms are analyzed and discussed in a related publication where the ability of this multispecies consortium to sustain hydrogen production is highlighted (Fakhimi et al., 2023b).

## TECHNICAL PROCEDURES

### Isolation of Stenotrophomonas goyi sp. nov

*S. goyi* sp. nov. was isolated from a fortuitously contaminated *Chlamydomonas reinhardtii* culture. Contamination occurred within the laboratory facilities (located at Campus Universitario de Rabanales, Cordoba, Spain). Initially, the *Chlamydomonas reinhardtii* culture was simultaneously contaminated with three different bacteria (Fakhimi et al., 2023b). Individual members of this bacterial community were isolated by sequential rounds of plate streaking in Yeast Extract Mannitol (YEM) medium, until 3 different types of bacterial colonies were visually identified. Colonies were grown separately, and the subsequent isolated DNA was used for PCR-amplification of their partial RNA 16S sequences. After sequencing, the three independently isolated bacteria were identified as members of the genus *Microbacterium, Stenotrophomonas, and Bacillus* (Fakhimi et al., 2023b).

### Genome sequencing and assembling of S. goyi sp. Nov

DNA isolated from and whole genome sequencing using PacBio (Pacific Biosciences) were performed by SNPsaurus LLC (https://www.snpsaurus.com/). Whole genome sequencing generated 102,238 reads yielding 832,209,774 bases for 166 read depth over the genome (Table 1). Genome was assembled with Canu 1.7 (Koren et al., 2017) yielding a 4,487,389 pb circular genome. The genome completeness was checked by Busco (Manni et al., 2021) and was 94.6% complete, with 94.6% of the genome single copy and 0.0% duplicated. Any other prokaryotic contamination was discarded using ContEst16S (Lee et al., 2017).

**Table 1.** Main de novo sequencing and assembly statistics.

| **Genome**<br>**Size (pb)** | **Fold-**<br>**coverage** | **GC content**<br>**(%)** | **Nº of**<br>**contigs** | **Type** | **Plasmids** |
| --- | --- | --- | --- | --- | --- |
| 4,487,389 | 166 | 66.5 | 1 | circular | No |

### Phylogenetic analysis

Phylogenetic analyses were performed using the TYGS (Type (Strain) Genome Server) (https://tygs.dsmz.de/) at Leibniz Institute DSMZ (German Collection of Microorganisms and Cell Cultures GmbH) (Meier-Kolthoff et al., 2022; Meier-Kolthoff and Göker, 2019). Information on nomenclature, synonymy and associated taxonomic literature was provided by TYGS’s sister database, the List of Prokaryotic names with Standing in Nomenclature (LPSN, available at https://lpsn.dsmz.de). The TYGS analysis was subdivided into 4 steps: 1) <u>Determination of closely related type strains</u>: Determination of closest type strain genomes was done in two complementary ways: First, genome was compared against all type strain genomes available in the TYGS database via the MASH algorithm (Ondov et al., 2016), and the ten type strains with the smallest MASH distances were chosen. Second, an additional set of ten closely related type strains was determined via the 16S rDNA gene sequences. These were extracted from the genome using RNAmmer (Lagesen et al., 2007) and each sequence was subsequently BLASTed (Camacho et al., 2009) against the 16S rDNA gene sequence of each of the currently 17208 type strains available in the TYGS database. This was used as a proxy to find the best 50 matching type strains (according to the bitscore) for the genome under study and to subsequently calculate precise distances using the Genome BLAST Distance Phylogeny approach (GBDP) under the algorithm ‘coverage’ and distance formula *d*_5_ (Meier-Kolthoff et al., 2013). These distances were finally used to determine the 10 closest type strain genomes for each of the user genomes. 2) <u>Pairwise comparison of genome sequences</u>: For the phylogenomic inference, all pairwise comparisons among the set of genomes were conducted using GBDP and accurate intergenomic distances inferred under the algorithm ‘trimming’ and distance formula *d*_5_ (Meier-Kolthoff et al., 2013). 100 distance replicates were calculated each. Digital DDH values and confidence intervals were calculated using the recommended settings of the GGDC 3.0 (Meier-Kolthoff et al., 2022, 2013). 3) <u>Phylogenetic inference</u>: The resulting intergenomic distances were used to infer a balanced minimum evolution tree with branch support via FASTME 2.1.6.1 including SPR postprocessing (Lefort et al., 2015). Branch support was inferred from 100 pseudo-bootstrap replicates each. The trees were rooted at the midpoint (Farris, 1972) and visualized with PhyD3 (Kreft et al., 2017). 4) <u>Type-based species and subspecies clustering</u>: The type-based species clustering using a 70% dDDH radius around each of the 12 type strains was done as previously described (Meier-Kolthoff and Göker, 2019). Subspecies clustering was done using a 79% dDDH threshold as previously introduced (Meier-Kolthoff et al., 2014). dDDH values were provided according to the following GBDP formulas: formula d0 (a.k.a. GGDC formula 1): length of all HSPs divided by total genome length; formula d4 (a.k.a. GGDC formula 2): sum of all identities found in HSPs divided by overall HSP length; formula d6 (a.k.a. GGDC formula 3): sum of all identities found in HSPs divided by total genome length. Note: Formula d4 is independent of genome length and is thus robust against the use of incomplete draft genomes.

### Annotation

The RAST tool kit, RASTtk at The Genome Annotation Service (Brettin et al., 2015; Overbeek et al., 2014) were used.

### Growth test and media

Bacterial precultures were grown on Yeast Extract Mannitol (YEM) or Lysogeny broth (LB) media. Some growth experiment were done in Mineral Medium (MM) (Harris, 2008) supplemented with different nutrient sources. Tris-Acetate-Phosphate (TAP) (Harris, 2008) were also used occasionally. More specific details for each experiment can be found in the corresponding figure and table captions. Bacterium cultures were incubated at 24-28°C and under continuous agitation (130 rpm).

### Coculturing algae and bacteria

Chlamydomonas cells were cultured for 3-4 days in TAP medium until mid-logarithmic growth phase, harvested by centrifugation (5.000 rpm for 5⍰min) and washed twice with fresh MM. Bacterial batch-cultures were incubated in TYM or LB medium until the Optical Density at 600⍰nm (OD_600_) reached 0.8-1, then harvested by centrifugation (12.000⍰rpm for 5⍰min) and washed twice with fresh MM. When needed a vitamin cocktail (riboflavin, 0.5 mg · L^-1^; p-aminobenzoic acid, 0.1 mg· L^-1^ ; nicotinic acid 0.1 mg· L^-1^ ; pantothenic acid, 0.1 mg· L^-1^ ; pyridoxine, 0.1 mg· L^-1^ ; biotin, 0.001 mg· L^-1^ ; vitamin B12, 0.001 mg· L^-1^ ; thiamine, 0.001 mg· L^-1^) was added to bacterial cultures. Algae and bacteria were cocultured in 250 mL flasks containing 100 mL of the corresponding medium. Alga and bacteria mixtures were set to reach initial chlorophyll concentration of 10 μg·mL^−1^ for the alga and an initial OD_600_ of 0.1 for the bacterium. Algal and bacterial monocultures were used as controls. All cultures were incubated at 24⍰°C with continuous agitation (80-140 rpm) and under continuous illumination (80 PPFD).

### Determinations of algal and bacterial growth

The algal growth was assessed in terms of chlorophyll content. Chlorophyll measurements were done by mixing 200⍰μL of the cultures with 800⍰μL of ethanol 100%. The mix was incubated at room temperature for 2-3 min, and afterward centrifuged for 1⍰min at 12.000⍰rpm. The supernatant was used to measure chlorophyll (a⍰+⍰b) spectrophotometrically (DU 800, Beckman Coulter) at 665 and 649⍰nm (Wintermans and de Mots, 1965).

Bacteria growth in monocultures was estimated spectrophotometrically in term of OD_600_ evolution (DU 800, Beckman Coulter). However, estimation of the bacterial growth in cocultures required bacterium cells separation from the alga cells. To do this, customized Selective Centrifugal Sedimentation (SCS) approach consisting in finding the centrifugation parameters that led to maximize algal cell sedimentation while minimize bacterial cell sedimentation was used (Torres et al., 2022). Measuring the O.D. of the supernatant after centrifugation can provide an estimation of the bacterial growth in the cocultures. To do this, the % of precipitated cells of each monoculture were calculated at different forces (from 100 x g to 500 x g) and times (1 and 2 min) using the measured O.D. before (A_BC_) and after (A_AC_) the centrifugation. By using 200 x g for 1 min led to 87.9% Chlamydomonas sedimentation, while only 2.1 % of the bacterial cells dropped (meaning that 97.9% of the *S. goyi* cells remained in the supernatant). This condition was chosen as a good compromise for SCS and used to evaluate the contribution of the bacteria to the OD in cocultures (^SCS^OD_600_).

### Data availability

*Stenotrophomonas goyi* sp. nov. has been deposited in the Spanish Type Culture Collection (CECT) with accession number CECT30764. The genome sequence for *Stenotrophomonas goyi* sp. nov. has been deposited as GenBank SUB12685123 at NCBI.

## RESULTS

### Identification of Stenotrophomonas goyi sp. nov

A fortuitous contaminated *Chlamydomonas reinhardtii* culture (strain 704) was studied due to its enhanced hydrogen production capability. This alga culture turned out to be contaminated with three different bacterial strain (Fakhimi et al., 2023b), one of them consisting in a white-pigmented bacterium (**Figure 1**). This bacterium was isolated after several rounds of plate streaking in TYM medium. First, partial PCR amplification and sequencing of the ribosomal 16S gene allowed the identification of this bacteria as a member of the *Stenotrophomonas* genus. Afterwards, the whole genome sequence was obtained. Genome assembling provided one single circular contig of 4,487,389 pb (**Table 1**). No plasmids or extrachromosomal elements were identified.

**Figure 1.**
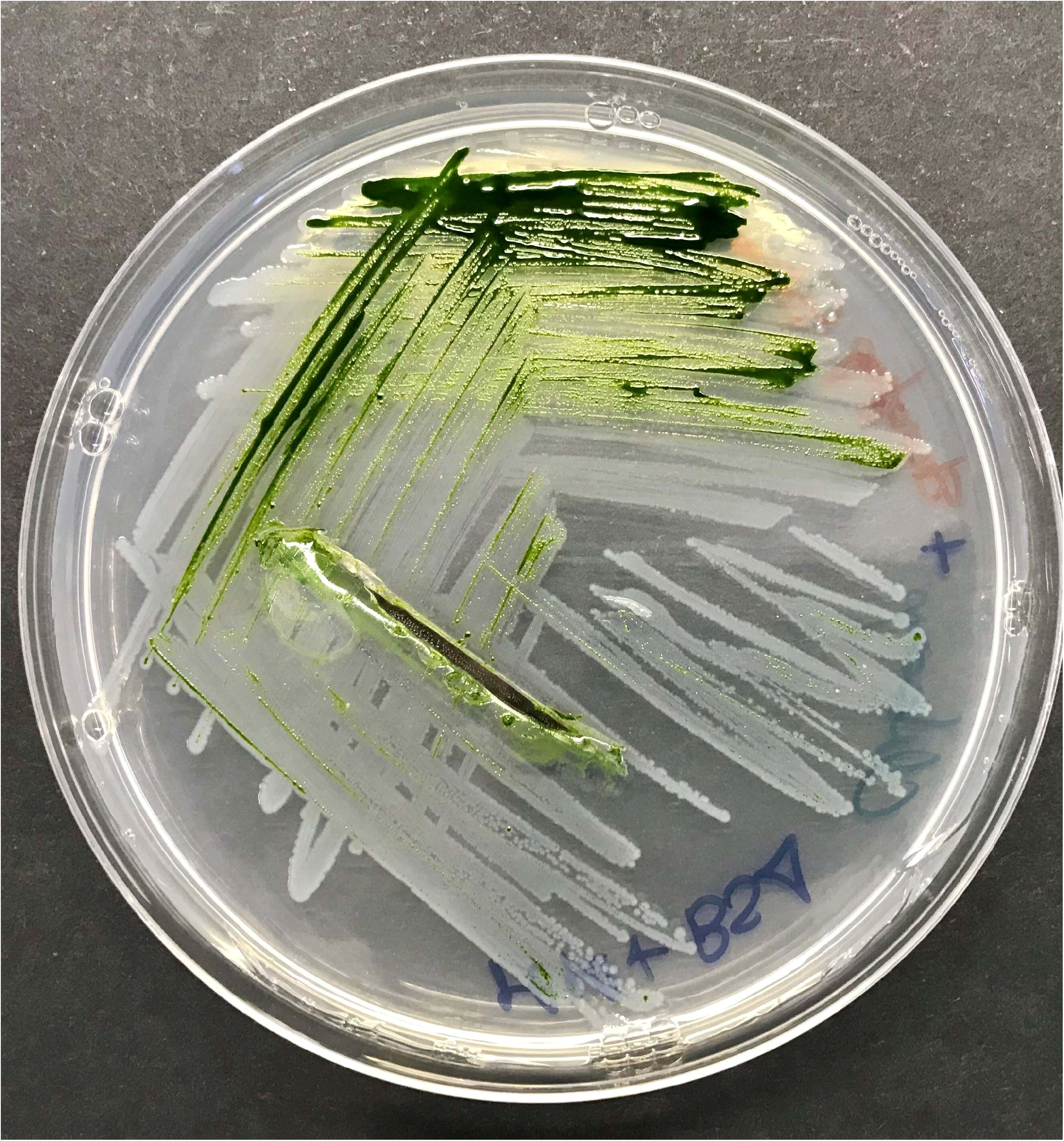
Plate with *Chlamydomonas reinhardtii* and *Stenotrophomonas goyi* sp. nov.

The RAST server identified 4,147 genes (4,066 CDS+ 81 rRNAs and tRNAs) (**Table 2**). Out of these 4,066 CDS identified by RAST, 1,096 of them were in subsystems. Tentative genome annotation derived from the RAST server is available in **Supplemental Table 1**.

**Table 2.** Genome features of *Stenotrophomonas goyi* sp. nov. according to RAST server.

| Genes | CDS | tRNAs | rRNAs |
| --- | --- | --- | --- |
| 4,147 | 4,066 | 71 | 10 |

Phylogenetic analyses were performed with both, the whole genome (**Fig. 2**) and the inferred 16S rDNAs (**Fig. 3**). Pairwise comparisons with the closest type strains genomes are shown in **Table 3**. These phylogenetic analyses revealed that the sequenced genome belonged to a new *Stenotrophomonas* sp.; all dDDH values (d0, d4 and d6) were below 70% (Meier-Kolthoff et al., 2013) (**Table 3**). This new bacterial specie was named as *Stenotrophomonas goyi* sp. nov. The closest related bacteria in terms of whole genome and 16S rDNA similarities were *Stenotrophomonas rhizophila DSM 14405 and Stenotrophomonas nematodicola W5*, respectively (**Fig. 2 and 3**). *S. goyi* sp. nov. genome was deposited in the NCBI as SUB12103906.

**Table 3.**
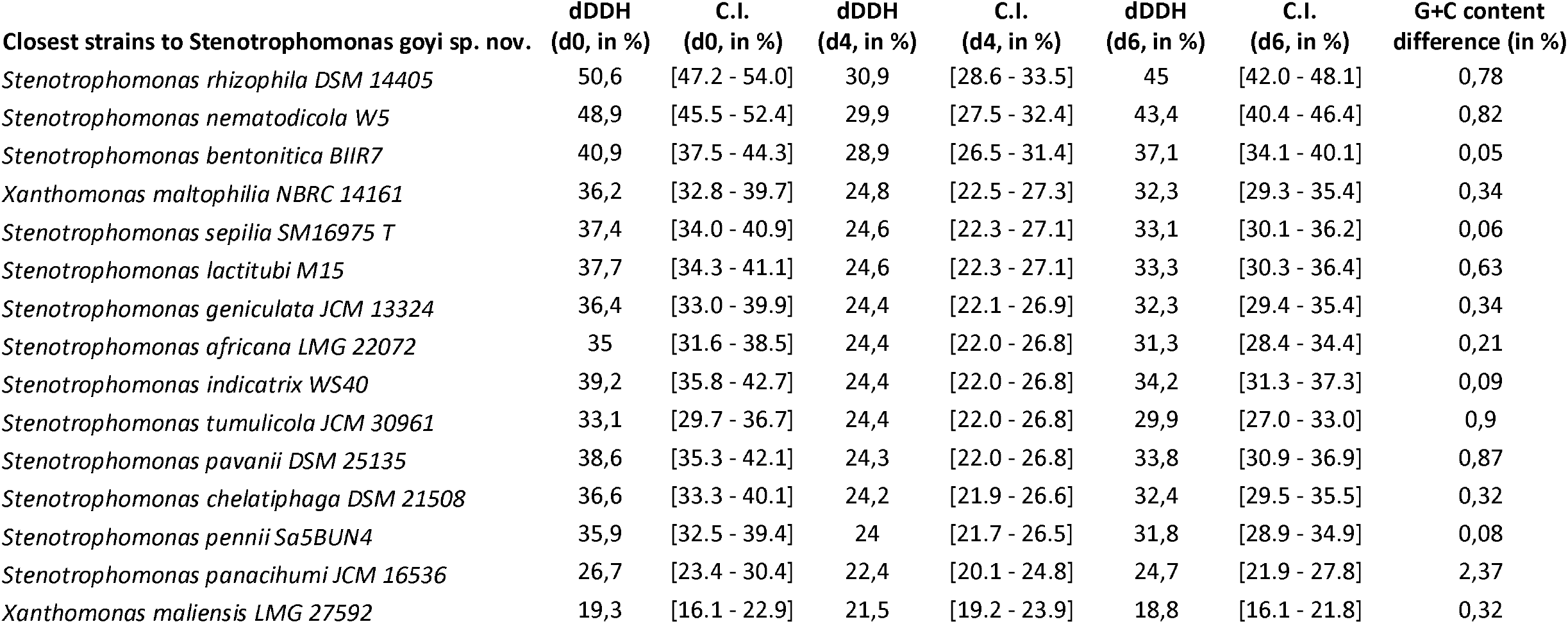
Pairwise dDDH values between S. goyi sp. nov and the closest type strains genomes. The dDDH values are provided along with their confidence intervals (C.I.) for the three different GBDP formulas: a) formula d0 (a.k.a. GGDC formula 1): length of all HSPs divided by total genome length; b) formula d4 (a.k.a. GGDC formula 2): sum of all identities found in HSPs divided by overall HSP length; formula d6 (a.k.a. GGDC formula 3): sum of all identities found in HSPs divided by total genome length. Note: Formula d4 is independent of genome length and is thus robust against the use of incomplete draft genomes.

**Figure 2.**
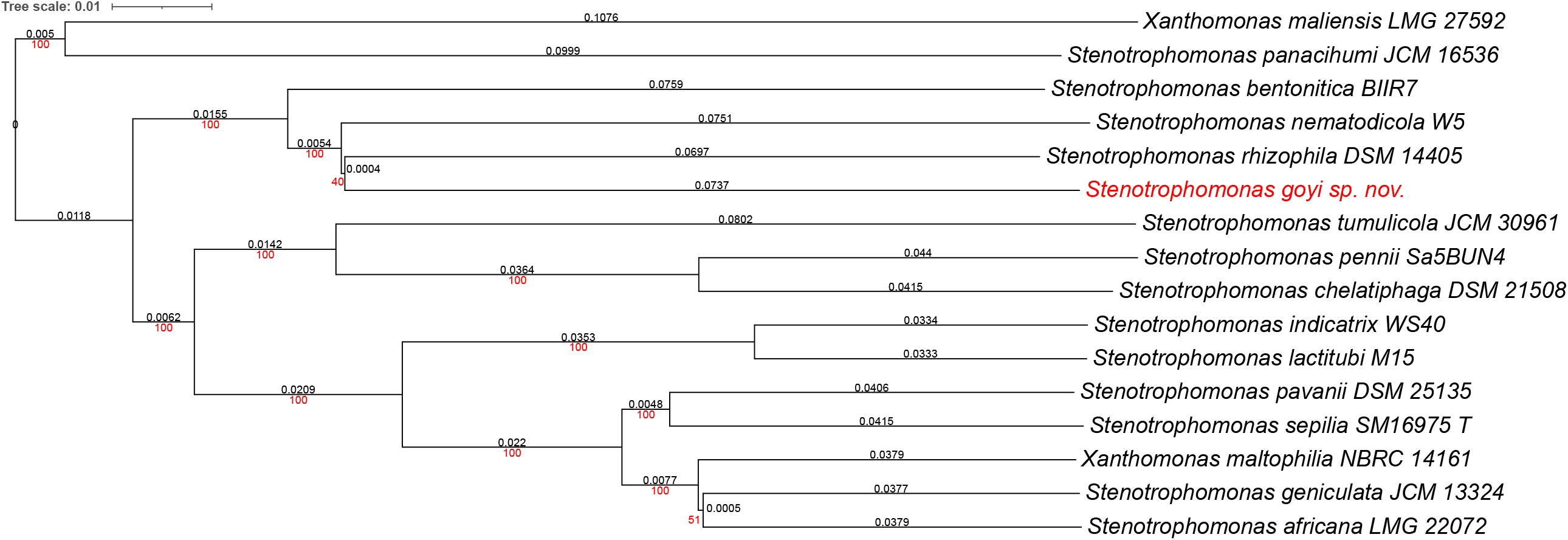
Phylogenetic tree for *Stenotrophomonas goyi* genome and related closest bacteria. Tree inferred with FastME 2.1.6.1 from GBDP distances calculated from genome sequences. The branch lengths are scaled in terms of GBDP distance formula d5. The numbers above branches are GBDP pseudo-bootstrap support values > 60 % from 100 replications, with an average branch support of 91.6 %. The tree was rooted at the midpoint. Branch lengths (black) and bootstraps (red) values are indicated. Genome sizes: 3,906,271 - 5,177,426 pb. Average δ statistics: 0.078 (Holland et al., 2002). Phylogenetic tree drawn with iTOL (Letunic and Bork, 2021).

**Figure 3.**
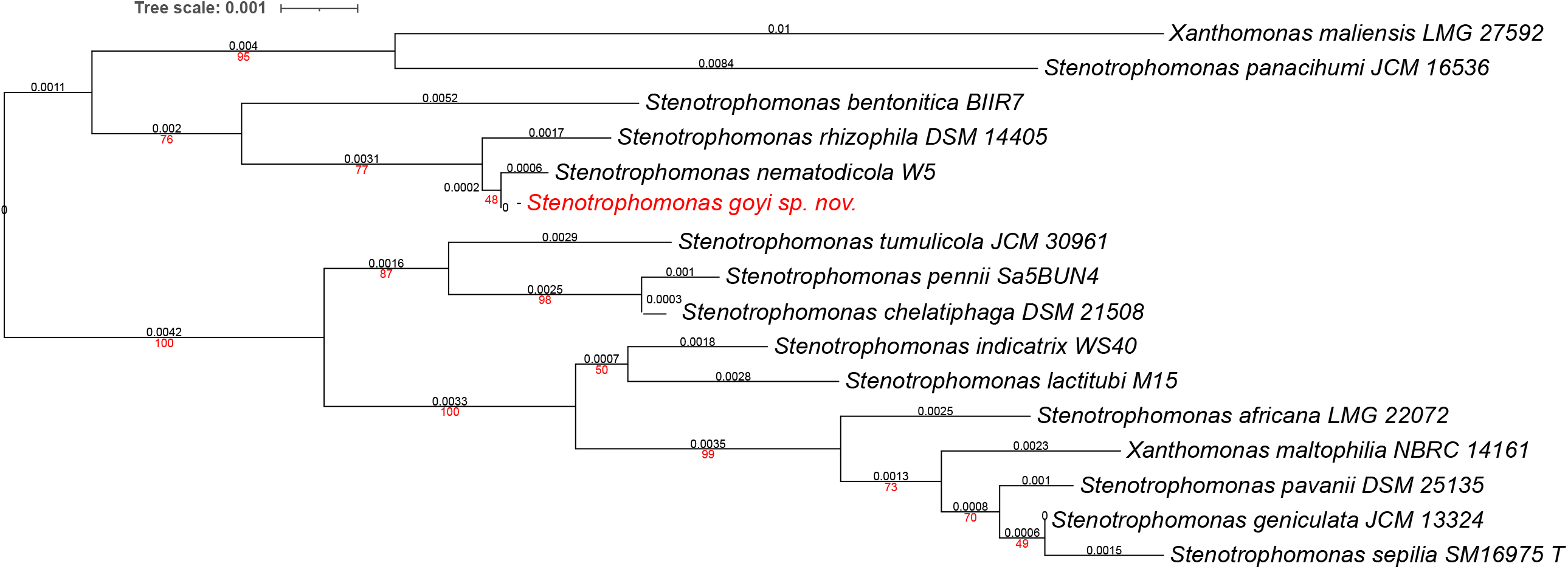
Phylogenetic tree for *Stenotrophomonas goyi* 16S rDNA and related closest bacteria. Tree inferred with FastME 2.1.6.1 from GBDP distances calculated from 16S rDNA gene sequences. The branch lengths are scaled in terms of GBDP distance formula d5. The numbers above branches are GBDP pseudo-bootstrap support values > 60 % from 100 replications, with an average branch support of 78.6 %. The tree was rooted at the midpoint. Branch lengths (black) and bootstraps (red) values are indicated. RNA16S lengths: 1,385 - 1,535 pb. Average δ statistics: 0.236 (Holland et al., 2002). Phylogenetic tree drawn with iTOL (Letunic and Bork, 2021)

BlastKOALA (Kanehisa et al., 2016) service allowed KEGG orthology assignments to characterize individual gene functions and reconstruct KEGG pathways of *S. goyi* genome (**Supplemental Table 2**). Some important pathways were either absent or incomplete in *S. goyi* sp. nov. including assimilation of nitrate (the whole assimilatory pathway is missing including nitrate transporters) and sulfate (only sulfite reductase is present). On the other hand, putative complete pathways for the glyoxylate cycle and biosynthesis of biotin, coenzyme A, pantothenate, riboflavin, tetrahydrofolate, glutathione, pyridoxal-P, lipoic acid, dTDP-L-rhamnose, UDP-N-acetyl-D-glucosamine, C5 isoprenoids, bacterial lipopolysaccharides, and antimicrobial proteins, among others, were present. Incomplete pathways for the degradation of aromatic compounds (including phenol, toluene, xylene, methylnafthalene, 3-hydroxytoluene, and terephthalate) myo-inositol biosynthesis, were also present.

Search with PHASTER (Arndt et al., 2016) revealed 1 intact prophage (PHAGE_Erwini_phiEt88_NC_015295) located at position 753112-799783 of the *S. goyi* sp. nov. genome.

### Nutrient requirements of S. goyi sp. nov

*S. goyi* sp. nov. showed no growth on MM, or in MM supplemented with different C sources (sucrose, glucose, lactose, mannitol, or glycerol) (**Table 4**). The addition of vitamins to MM supplemented with glucose or lactose also did not support the bacterium growth (**Table 4**). However, the bacterium showed an excellent growth when cultivated in MM supplemented with yeast extract, tryptone, peptone or even BSA (**Table 4**), suggesting that this bacterium has a great capacity to use peptides/amino acids as C source, and probably also as N source. Moreover, the peptides/amino acids could also provide, in addition to C and N sources, with other essential nutrients or even palliate potential amino acids auxotrophies. Note that MM medium has sulfate as only S source. As commented before, the genome of *S. goyi* sp. nov. is lacking a functional sulfate assimilation pathway. Thereby S-containing amino acids (cysteine and methionine) could support the growth in medium rich in peptides/amino acids.

**Table 4.** Growth of *S. goyi* sp. nov. with different carbon sources. MM medium was supplemented with different nutrients at 5 g·L^-1^ each, but methanol and ethanol (5 mL·L^-1^). For acetic acid, TAP medium was employed (1.05 g·L^-1^ of acetic acid). Vitamins cocktail included riboflavin (0.5 mg· L ^-1^), p-aminobenzoic acid (0.1 mg· L ^-1^), nicotinic acid (0.1 mg·L^-1^), pantothenic acid (0.1 mg· L ^-1^), pyridoxine (0.1 mg· L ^-1^), biotin (0.001 mg· L ^-1^), vitamin B12 (0.001 mg · L^-1^), thiamine (0.001 mg· L ^-1^). ++, significant growth; +, poor growth; -, no growth.

|  | Growth |
| --- | --- |
| Sucrose | - |
| Lactose | - |
| Lactose + vitamins | - |
| Glucose | - |
| Glucose + vitamins | - |
| Mannitol | - |
| Lactic acid | - |
| Glycerol | - |
| Acetic acid (TAP medium) | - |
| Tryptone | ++ |
| Peptone | ++ |
| Yeast extract | ++ |
| BSA | ++ |

To confirm this hypothesis, *S. goyi* sp. nov. was inoculated in plates of MM medium + glucose supplemented with different combinations of cysteine, methionine, biotin, and thiamine. Only plates containing cysteine and methionine supported the bacterial growth for several culturing rounds (**Fig. 4**). This result confirms the cysteine and methionine growth dependence of *S. goyi*. sp. nov. Cysteine and methionine could provide with either a S-reduced source or complement an auxotrophy for these two amino acids. Since *S. goyi* sp. nov. genome has complete pathways for all the essential amino acids, is more likely that cysteine and methionine are being used as S-reduced sources. Similar results were found for *M. fakhimi*, where cysteine and methionine are required as S-sources (Fakhimi et al., 2023a).

**Figure 4.**
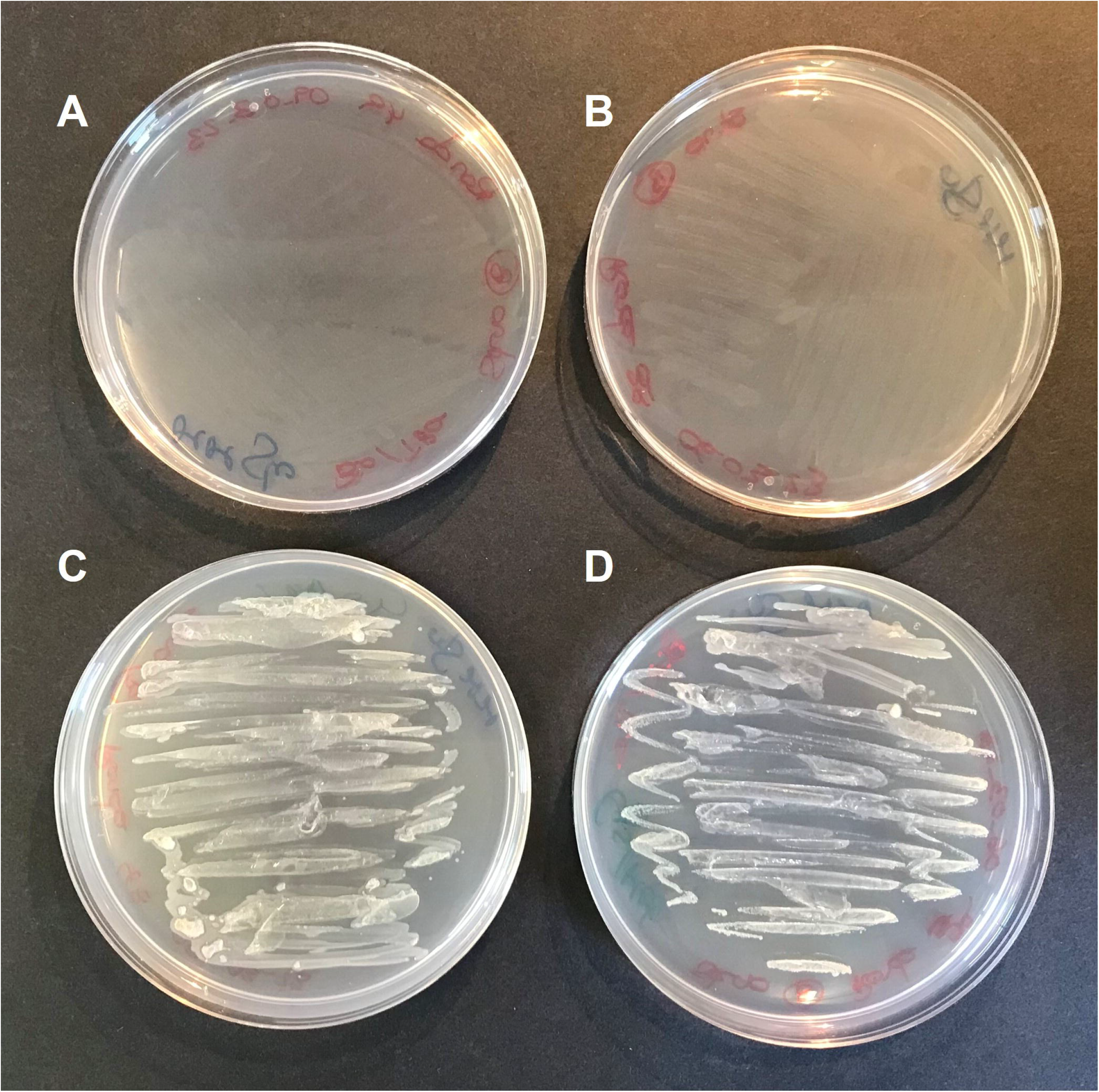
Cysteine and methionine requirements of S. goyi to grow. *S. goyi* sp. nov. was inoculated in: A) plates of MM medium + glucose (5 g·L^-1^); B) MM medium + glucose + biotin (0.001 mg·L^-1^) + thiamine (0.001 mg·L^-1^); C) MM medium + glucose + cysteine (4 mM) + methionine (4 mM); and D) MM medium + glucose + cysteine + methionine + biotin + thiamine.

*S. goyi* sp. nov. showed optimal growth between 24<u>^°^</u>C and 32<u>^°^</u>C and pH 5-9 (**Table 5**). Despite the presence in the genome of the complete multidrug resistance efflux pump MexJK-OprM (Chuanchuen et al., 2005), no resistance to the antibiotics tetracycline, rifampicin, chloramphenicol and polymyxin (50 μg/mL each) was observed (data not shown).

**Table 5.**
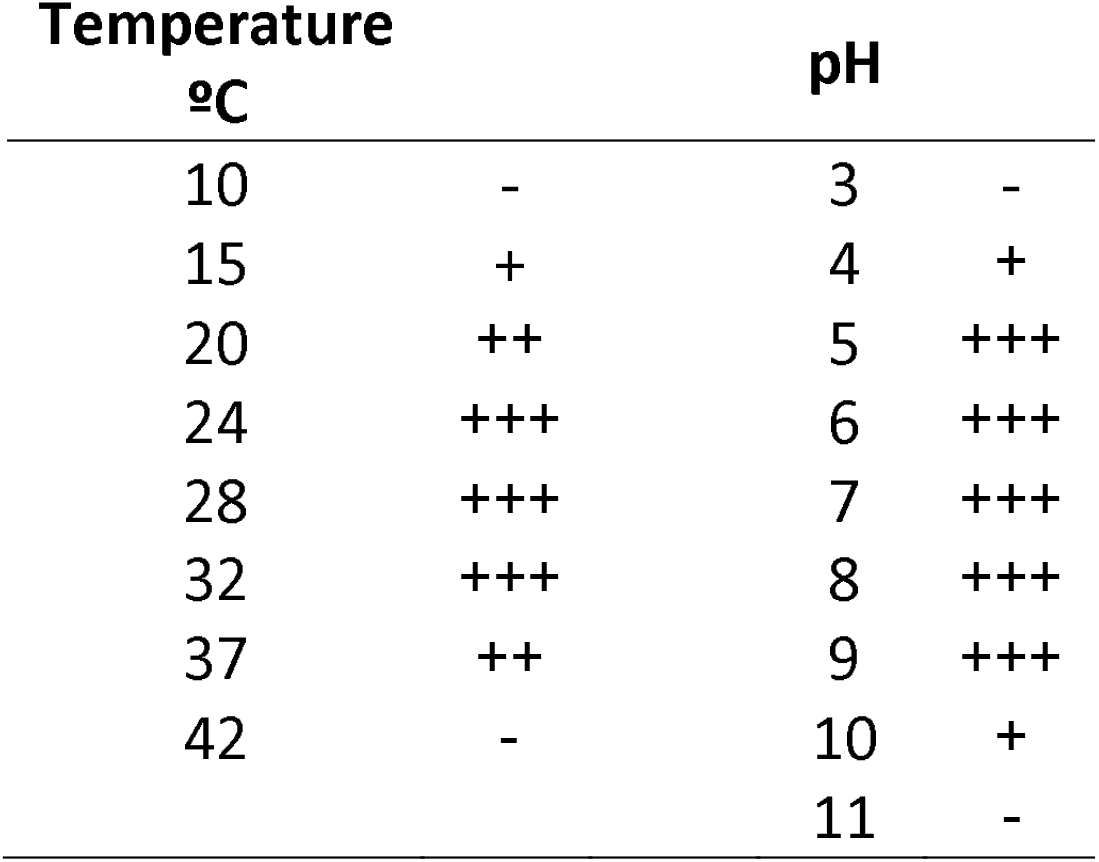
*S. goyi* sp. nov. growth at different temperatures and pH. LB medium was used in all the experiments.

### Growth of S. goyi sp. nov -C. reinhardtii consortium

Torres et al., (2022) reported that cocultures of *S. goyi* sp. nov (published as *Stenotrophomonas* sp.) and *C. reinhardtii* promoted the growth of the microalga (nearly doubled) when incubated in MM supplemented with glucose and mannitol, but not when supplemented with acetic acid (Torres et al., 2022).

Here, we checked if the bacterium also benefited when cultivated in glucose and mannitol containing media in the consortium with C. *reinhardtii*. First, we observed that the chlorophyll content after 13 days of coculturing was 2.4 times higher than the *C. reinhardtii* monocultures (**Fig. 5a**), which is in accordance with previous results (Torres et al., 2022). Additionally, the dry biomass resulting from the consortia was 2.2 times higher than the sum of the respective monocultures (**Fig. 5b**). Finally, the growth of the bacterium in cocultures (estimated using a SCS approach (Torres et al., 2022)) showed that *S. goyi* sp. nov. grew very efficiently, unlike *S. goyi* sp. nov. monocultures (**Fig. 5c**).

**Figure 5.**
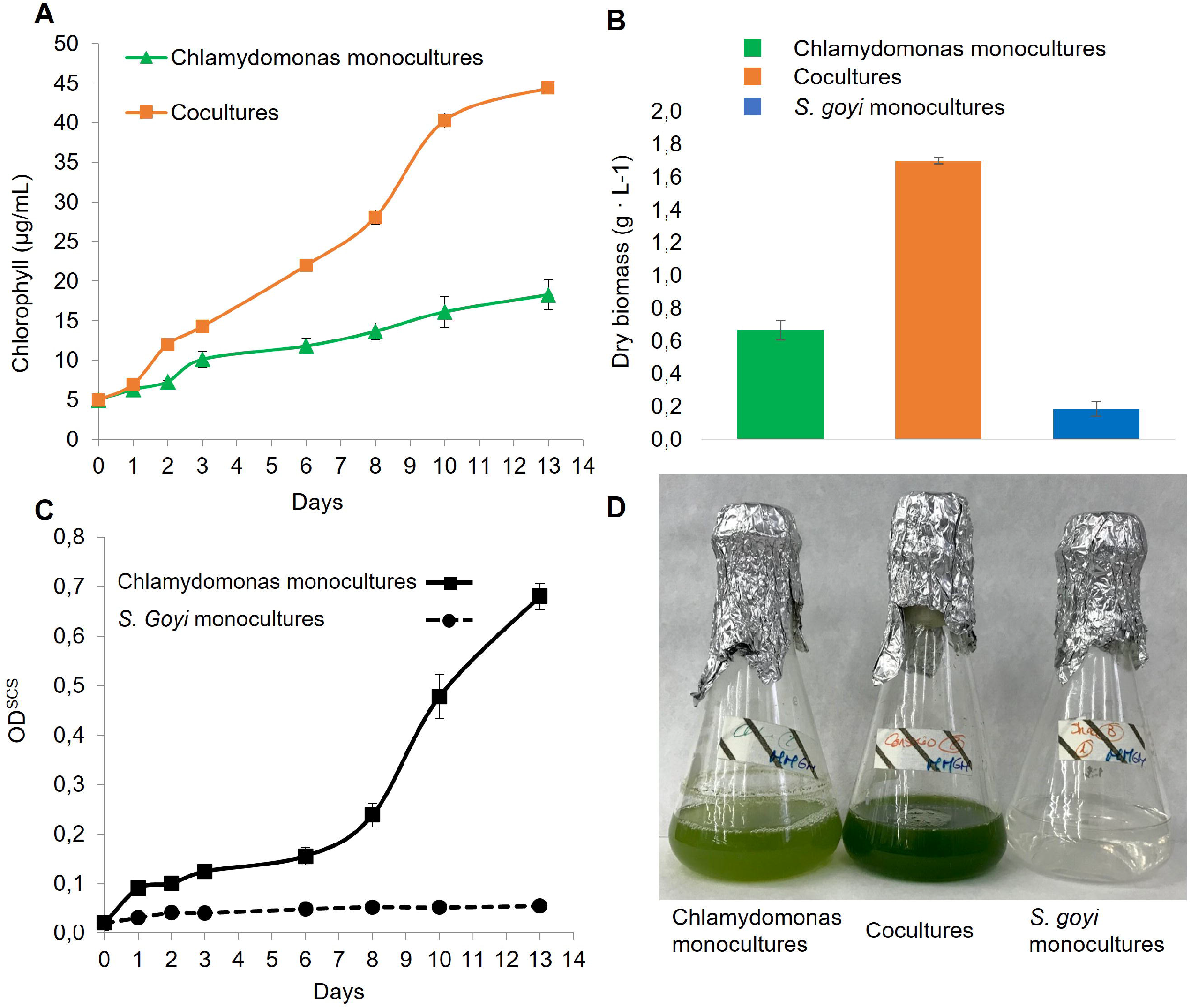
*S. goyi* sp. nov. and *C. reinhardtii* growth in consortium. *S. goyi-C. reinhardtii* consortium, and respective control monocultures, were incubated in MM supplemented with glucose (5 g·L^-1^) and mannitol (5 g·L^-1^). A) Chlorophyll content; B) Dry weight biomass after 13 days; C) OD^SCS^; D) Actual picture of the cultures after 13 days.

These results indicate that *S. goyi* sp. nov. and *C. reinhardtii* can establish a mutualistic relationship when incubated in sugars-containing media. On one hand, *S. goyi* sp. nov. can greatly support the growth of the *C. reinhardtii* in media supplemented with glucose or mannitol, which are two carbon sources that the alga cannot utilize. Likely, this growth promotion is due to the release of acetate and/or CO_2_ from the bacteria after the sugar fermentation. Acetate is the solely organic carbon source that *C. reinhardtii* can utilize during heterotrophic/mixotrophic growth (Chaiboonchoe et al., 2014). On the other hand, *S. goyi*, sp. nov. can grow in media without amino acids/peptides supplementation if cocultured with *C. reinhardtii*, suggesting that the alga must provide some essential nutrients for the bacterium. S-reduced forms excreted by the alga (e.g., cysteine or methionine) could potentially explain the bacterium growth in the consortium.

### Biotechnological importance of Stenotrophomonas goyi sp. nov. in the microalga context

Stenotrophomonas spp. are common constituent of the rhizosphere, and their potential for agricultural biotechnology is arising. However, their association with algae is poorly explored. Most plant growth-promoting bacteria (PGPB) are firstly identified in the rhizosphere and in association with plants. However, many PGPB are then also often commonly found in association with algae. Probably this is reflecting that the kind of relationships established between bacteria and plants are similar to the relationships between bacteria and algae. This could be potentially the case of Stenotrophomonas spp., although the relatively youth and heterogeneity of the genus may prevent tracking down their association with algae.

Some Stenotrophomonas spp. show a limited nutritional range while some others are capable of metabolic versatility (Ryan et al., 2009). *S. goyi* sp. nov. is unable to grow in the absence of a source of peptides/amino acids, which imply that in natural ecosystems it may rely on other microorganisms to obtain essential nutrients. This requirement for peptides/amino acids is likely related to the obtaining of S-reduced forms through cysteine and methionine since *S. goyi* sp. nov. is unable to uptake sulfate.

*Stenotrophomonas goyi* sp. nov. was isolated from an alga culture (*C. reinhardtii*) that showed an enhanced capacity to produce hydrogen when incubated in medium with mannitol and yeast extract (Fakhimi et al., 2023b). This algal culture was simultaneously contaminated with two other bacteria: *Microbacterium fakhimi* and *Bacillus cereus*. Although the main responsible of the enhance algal hydrogen production observed in this contaminated alga culture was *M. fakhimi* sp. nov., the *C. reinhardtii-M. fakhimi* cocultures were unable to produce hydrogen and biomass concomitantly. However, when the four microorganisms were cultivated together (C. *reinhardtii, S. goyi, M. fakhimi* and *B. cereus*) the simultaneous obtaining of both hydrogen and biomass was possible (Fakhimi et al., 2023b), which extend the biotechnological interest.

In this multispecies association, *S. goyi* and *C. reinhardtii* could be alleviating the auxotrophy of *M. fakhimi* sp. nov. for biotin and thiamine. S. goyi sp. nov. could also provide ammonium derived from the mineralization of the amino acids for the alga. On the other hand, the alga could provide S-reduced sources such as cysteine and methionine for *S. goyi* sp. nov. In any case, this multispecies association was mutually beneficial and prevented an excessive bacterial growth in cocultures, which could be one of the main drawbacks when algae-bacteria cocultures are used for biotechnological applications.

However, the precise metabolic relationships established in this multispecies consortium that led to extend the viability of the *C. reinhardtii* cells during hydrogen production conditions is not unraveled yet and need to be further investigated.

## Author Contributions

Writing—original draft preparation, D.G.-B.; writing—review & editing, N.F. A.D.; Isolation of the *Stenotrophomonas goyi* sp. nov., N.F.; design and execution of experiments, M.J.T; in silico analysis, D.G.-B.; results analysis and interpretation, M.J.T, N.F., D.G.-B. and A.D.; supervision, A.D. and D.G.-B.; project administration, A.D. and D.G.-B.; funding acquisition, A.D. and D.G.-B. All authors have read and agreed to the published version of the manuscript.

## Acknowledgments

The authors acknowledge Dr. Gregorio Galvéz Valdivieso for his unvaluable contribution to this research: perfection sometimes kills new discoveries. We thank the Bio knowledge Lab (BK-L) Ltd. for their kind support.

## Figure legends

**Table 1. Main de novo sequencing and assembly statistics of *Stenotrophomonas goyi* sp. nov. genome**

**Table 2. Genome features of *Stenotrophomonas goyi* sp. nov. according to the RAST server**

**Table 3. Pairwise dDDH values between *S. goyi* sp. nov. and the closest type strains genomes**. The dDDH values are provided along with their confidence intervals (C.I.) for the three different GBDP formulas: a) formula d0 (a.k.a. GGDC formula 1): length of all HSPs divided by total genome length; b) formula d4 (a.k.a. GGDC formula 2): sum of all identities found in HSPs divided by overall HSP length; formula d6 (a.k.a. GGDC formula 3): sum of all identities found in HSPs divided by total genome length. Note: Formula d4 is independent of genome length and is thus robust against the use of incomplete draft genomes.

**Table 4. Growth of S. goyi sp. nov. on different nutrients**. MM medium was supplemented with different nutrients at 5 g·L^-1^ each, but methanol and ethanol (5 ml·L^-1^). For acetic acid, TAP L), medium was employed (1.05 g·L^-1^ of acetic acid). Vitamins cocktail included riboflavin (0.5 mg· L^-1^ p-aminobenzoic acid (0.1 mg· L ^-1^), nicotinic acid (0.1 mg· L ^-1^), pantothenic acid (0.1 mg· L ^-1^), pyridoxine (0.1 mg· L ^-1^), biotin (0.001 mg· L ^-1^), vitamin B12 (0.001 mg· L ^-1^), thiamine (0.001 mg· L ^-1^). ++, significant growth; +, poor growth; -, no growth.

**Table 5. Growth of *S. goyi* sp. nov. at different temperatures and pHs**. LB medium was used in all the conditions.

**Supplemental Table 1. Tentative genome annotation of S. goyi sp. nov**.

**Supplemental Table 2. BlastKOALA gene orthology assignments**

